# Preclinical establishment of a divalent vaccine against SARS-CoV-2

**DOI:** 10.1101/2022.02.10.479919

**Authors:** Zsofia Hevesi, Daniela Gerges, Sebastian Kapps, Raimundo Freire, Sophie Schmidt, Daniela D. Pollak, Klaus Schmetterer, Tobias Frey, Rita Lang, Wolfgang Winnicki, Alice Schmidt, Tibor Harkany, Ludwig Wagner

**Author notes:** **Correspondence:** Dr. Ludwig Wagner (at Division of Nephrology and Dialysis, Department of Medicine III, Medical University of Vienna, Vienna, Austria), *e-mail:. These authors contributed equally (Zsofia Hevesi, Daniela Gerges).

## Abstract

First-generation vaccines against SARS-CoV-2 have been administered to more than 60% of the population in developed countries. However, the monovalent vaccines currently available in Europe do not confer adequate and durable immune protection. To satisfy the need for a novel vaccine, we engineered a divalent gene construct consisting of the receptor binding domain (RBD, 300-685 aa) of the spike protein and the immunodominant region of the nucleocapsid (100-300 aa). This fusion protein was cloned into a pET-30a plasmid and expressed either in *Escherichia coli* or in a recombinant baculovirus in insect cells. Following purification *via* its His-tag, the fusion protein was mixed with adjuvant, and administered to mice in a prime-booster-mode. Upon testing for IgG antibody response against nucleocapsid and RBD, a titer of 10^−4^ - 10^−5^ was demonstrated 14 days after the first booster injection in 72% of the animals, which could be increased to 100% by a second booster. Notably, comparable IgG responses were detected against the delta, gamma and omicron variants of the RBD region. Durability testing revealed the presence of IgG beyond 90 days. In addition, granzyme A and perforin mRNA expression (cytolytic effector cell molecules) was increased in cytotoxic lymphocytes isolated from peripheral blood. *Ex vivo* stimulation of T-cells by nucleocapsid and RBD peptides showed antigen-specific upregulation of CD44 in vaccinated mice among their CD4^+^ and CD8^+^ T-cells. No side-effect was documented in the central nervous system, be it either endothelial inflammation or neuronal damage. Cumulatively, the combined induction of B-cell and T-cell response by a bivalent protein-based vaccine directed against two structural SARS-CoV-2 proteins represents a proof-of-principle approach alternative to existing mRNA vaccination strategies, which could confer long-lasting immunity against all known viral strains.

## Introduction

Many efforts have been made to generate spike glycoprotein-based vaccines to induce immunoprotection against SARS-CoV-2 infection. Both the mRNA-based (Jackson, Anderson et al., 2020, Mulligan, Lyke et al., 2020) and protein-based (Guebre-Xabier, Patel et al., 2020) vaccines are effective in humans. Further protein-based designs at a preclinical stage of development promise superior safety and efficacy, at least in animal models (Watterson, Wijesundara et al., 2021). However, there has been and continues to be a growing concern that existing and emerging virus mutants may escape currently-available vaccine-induced immunity (Mostaghimi, Valdez et al., 2021). Although vaccinated individuals infected with the delta or omicron variants of SARS-CoV-2 have a lower infectious viral titer (IVT) than non-vaccinated individuals (Puhach, Adea et al., 2022), the spread of these variants is very rapid, despite high rates of vaccination using mRNA or vector vaccines encoding the spike protein alone. Therefore, proposals have been made to include other SARS-CoV-2 proteins known to generate high antibody titers and T-cell responses (Ahlen, Frelin et al., 2020, Dutta, Mazumdar et al., 2020, Li, Wang et al., 2020).

One of the four structural proteins ubiquitously required for viral replication is the nucleocapsid, which is immediately translated and expressed at excess quantities so that it is even secreted from infected cells into the blood. As a result, nucleocapsid detection is an effective means to screen infected individuals even before the onset of symptoms (Diao, Wen et al., 2020, Li et al., 2020), particularly because antibody production against this protein is rapid and abundant (Burbelo, Riedo et al., 2020, Guo, Ren et al., 2020). The 100-300 aa region of the nucleocapsid induces the highest antibody titer in humans (Smits, Hernández-Carralero et al., 2021), which is attributed to its octapeptide structure homologous to the four hyperendemic seasonal coronaviruses (Pajenda, Kapps et al., 2021). Epitope profiling of SARS-CoV-2 antibodies with cross-reactivity to seasonal coronaviruses (Ladner, Henson et al., 2021) together with experimental data showing that an antibody against the nucleocapsid can protect mice from lethal infection with a hepatitis virus (Nakanaga, Yamanouchi et al., 1986) highlight the conceptual and technical potential of targeting this protein epitope. In addition, recent work in hamsters showed that immunization with attenuated intracellular replicating bacterium-vectored vaccines protects from the typical disease pathology caused by SARS-CoV-2 infection (Jia, Bielefeldt-Ohmann et al., 2021). Alternatively, specific antibodies raised against the receptor binding domain (RBD) of the spike protein can neutralize and inhibit the entry of SARS-CoV-2 into angiotensin-converting enzyme-2-expressing cells (Wu, Wang et al., 2020). As such, the RBD is an immunodominant and specific target of SARS-CoV-2 antibody production in virus-infected individuals (Premkumar, Segovia-Chumbez et al., 2020).

Here, we generated a fusion protein that combines the nucleocapsid (*<u>N</u>*100-300 aa) and the RBD of the spike protein (*<u>S</u>*300-685 aa) in one molecule. The recombinant construct was introduced into bacterial and insect expression vectors and produced in *Escherichia coli* (*E. coli*) and insect cells, respectively. Its immunogenic efficacy was shown by murine vaccination studies using a prime-booster strategy, tested against all strains of SARS-CoV-2, provided prolonged immune responses and lacked any appreciable side-effect, particularly those occurring in the brain and related to mortality in humans exposed to other vaccines (Greinacher, Thiele et al., 2021, Rzymski, Perek et al., 2021, Zuhorn, Graf et al., 2021).

## Methods

### Animals, blood sampling and tissue processing

A total of 18 male and 4 female mice (C57BL/6J, 8–12-week-old) were group housed under standard conditions with a 12/12 light/dark cycle. The Austrian Federal Ministry of Education, Science and Research granted approval for this research (2021-0.721.018). Experimental procedures were approved by the Austrian Ministry of Science and Research (66.009/0145-WF/II/3b/2014 and 66.009/0277-WF/V/3b/2017) and conformed to the 2010/63 European Communities Council Directive. Mice were habituated for at least a week to their environments and their numbers were kept at an absolute minimum. Blood was collected from the facial vein at a maximum volume of 200 µl every other week. At the end of the post-immunization survival period, mice were deeply anesthetized by isoflurane (at 5% with 1 L/min flow rate of tubed air) and then perfusion-fixed by transcardially applying 4% (wt/vol) paraformaldehyde and 0.1% glutaraldehyde in 0.1 M phosphate buffer (PB; pH 7.0). Dissected brains were immersed in the same fixative (without glutaraldehyde) at 4 °C overnight. Brains were cryoprotected in 30% sucrose in PB at 4 °C for 3 days. Coronal sections (50 μm) were cut on a cryostat microtome (1-in-4 series) and kept in 0.05% NaN_3_ in PB until immunohistochemical processing.

### Construction and heterologous expression of the fusion protein

The fusion protein was constructed using the Gibson assembly method (Gibson, Young et al., 2009). For vector construction, the portion of the nucleocapsid (*<u>N</u>*100-300 aa) fused to the RBD (*<u>S</u>*300-685 aa) including 4 glycines as a hinge region was cloned into a pET-30a vector and designated ‘VieVac’ (**Supplementary Figure 1A**). This product was generated by first producing 2 fragments by PCR using the *N* and *S* cDNAs obtained from the Krogan laboratory as template (Gordon, Jang et al., 2020). The fragment containing the complete *N* protein protion and the beginning of the *S* protein were amplified with the primers as follows: forward ATGGCTGATATCGGATCCGAATTCATGAAAGATCTCAGTCCGCGCTGG and reverse TTTAAGTGTACAACCACCGCCACCATGTTTGTAATCTGTCCCTTGCCG. To generate the second overlapping fragment containing the end of the *N* protein and the entire *S*-RBD sequence, we used the following primers: forward GATTACAAACATGGTGGCGGAGGTTGTACACTTAAAAGTTTTACGGTC and reverse GCCGCAAGCTTGTCGACGGAGCTCTCATCGCGCTCTTCGCGGGGAATT. In the final cloning step, the EcoR1 linearized pET-30a plasmid was combined with fragments 1 and 2.

### Directional cloning into the Gateway pEntry/D-TOPO vector

A 4-nucleotide overhang (CACC) was placed in front of the forward primer CACCATGAAAGATCTCAGTCCGCG, while TCATCGCGCTCTTCGCGGGG served as reverse primer. The original construct (engineered pET-30a/VieVac) has been used as template. The PCR conditions were as: 95°C for denaturing, 58°C for 30 sec annealing and 72°C for 2 min synthesis with 30 cycles. The resultant PCR amplicon was ligated into the pEntry/D-TOPO vector by incubation at 24°C for 1h. The resultant plasmid was transfected into One Shot TOP10 chemically competent *E. coli* using the heat shock method. Following 60 min incubation at 37 °C, *E. coli* were spread onto LB/kanamycin plates and incubated at 37 °C overnight. Outgrowing colonies were amplified in 3 ml LB/kanamycin in a shaking incubator (200/min) for 14 h. Plasmid isolation was carried out using the PureLink™ Quick Plasmid Miniprep Kit (Invitrogen).

### Insertion into baculovirus and amplification in insect cells

The pEntry/D-TOPO/VieVac construct (25 ng) was shifted into the baculovirus-compatible pDEST™ 10 expression vector. Using Clonase LR-Reaction II, both plasmids (pEntry/D-TOPO/VieVac and pDEST™ 10) were combined together with 1 µl 5x LR-Clonase II enzyme in a final volume of 5 µl. This mixture was incubated at 25 °C for 3 h. One µl was used for transforming One Shot chemically competent *E. coli* by applying the heat shock method. Following 60 min incubation at 37 °C, the cells were spread onto LB/ampicillin agar plates and incubated at 37 °C overnight. The outgrowing clones were amplified in liquid culture (LB/ampicillin) with plasmids isolated using the PureLink Quick Plasmid Miniprep Kit. Following sequence verification, the Bac-to-Bac® baculovirus system (Invitrogen) was chosen to generate the VieVac bacmid. In brief, Max Efficiency® DH10Bac™ competent *E. coli* were transformed with the engineered pDEST™ 10 containing the VieVac recombinant construct using the heat shock method at 42 °C for 30 s. Transformants were grown at 37 °C under shaking in SOC medium for 4 h. A 10-fold serial dilution of the cells was made with each of them spread for selection on 7 µg/ml gentamicin, 10 µg/ml tetracycline, 50 µg/ml kanamycine, 100 µg/ml BluoGal and 40 µg/ml IPTG-containing LB agar plates. White colonies appearing after 48 h were re-plated and incubated at 37 °C for 14 h. Colonies with white phenotype were grown out in liquid culture to isolate recombinant bacmid DNA by the Miniprep method. Following insert verification by PCR, Sf9 insect cells were transfected using Cellfectin® II reagent. Eight µl of Cellfectin® was diluted in 100 µl Grace’s Insect Medium (unsupplemented) and 1 µg of recombinant baculovirus DNA in 100 µl Grace’s Insect Medium (unsupplemented). We then mixed diluted Cellfectin® and baculovirus DNA, incubated these for 15 min at room temperature, and added to the insect cells freshly plated 1 h earlier. The transfection medium was replaced by protein expression medium (SFM4Insect™ with L-Glutamine). Protein expression was confirmed by a monoclonal antibody against the His-tag HIS.H8 (*see below*).

### ‘VieVac’ purification from insect cells

Seventy-two h after infection, Sf9 or Hi5 cells were centrifuged at 600 *g* for 5 min. Cells in the pellet were lysed using diluted BugBuster Protein Extraction Reagent® (1:3 in PBS). After lysate sonication for 10 s and addition of benzonase endonueclease, the insoluble material was pelleted at 12,000 *g* at 4 °C for 10 min. The pellet was re-solubilized in 6M GuHCl, 0.1M NaH_2_PO_4_, 0.01M Tris-Cl (pH 8.0) and passed through a Ni-NTA binding column (Qiagen). The loaded column was washed with 8M urea, 0.1M NaH_2_PO_4_, 0.01M Tris-Cl (pH 8,0) followed by 8M urea, 0.1M NaH_2_PO_4_, 0.01M Tris-Cl (pH 6.3) as also described for proteins produced in *E. coli* (*see below*). Finally, elution of the recombinant protein was carried out with 8M urea, 0.1M NaH_2_PO_4_, 0.01M Tris-Cl (pH 4.5) containing 250 mM imidazole. The resultant eluate was tested for protein content by spectrophotometry and immunoblotting.

### Evaluation of protein expression by immunofluorescence

In order to test ‘VieVac’-fusion protein expression in insect cells, Sf9 and Hi5-cells suspended in tissue culture medium were employed to generate cyto-slide preparations using a cyto-centrifuge. Air-dried cyto-preparations were fixed in acetone for 5 min. A water repellent circle was drawn around the area where cells were spread. Following wetting with phosphate buffered saline (PBS, pH 7.4), 60 µl of pre-diluted HIS.H8 mAb (Sigma) was applied onto the cells and incubated for 2 h under continuous agitation at room temperature in a moisturized chamber. After washing in PBS for 10 min, Alexa Fluor 594-tagged goat anti-mouse antibody was applied for 1 h at room temperature under continued gentle agitation. Hoechst 33,342 was routinely used as nuclear counterstain. Following repeated washes, specimens were mounted by Vectashield and coverslipped. Images were captured on a confocal microscope (Zeiss 880).

### Protein production and purification from *E. coli*

The engineered pET-30a/VieVac vector was transformed into BL21 (DE3) *E. coli* using the heat shock method: Two µl of plasmid were mixed with 100 µl BL21 (DE3) or BL21 Codon Plus *E. coli* and incubated on ice for 30 min. Following a 45 s heat shock at 42 °C, the samples were returned to ice for 5 min. SOC Medium was applied and bacteria were incubated for 60 min at 37 °C under constant shaking. Aliquots were spread onto LB/kanamycin plates, which were incubated at 37 °C overnight. Kanamycin-resistant colonies were expanded in liquid culture using LB/kanamycin broth overnight. The culture volume was expanded 5x with Terrific Broth and induced with 200 µmol isopropyl-ß-D-thiogalactopyranosid (IPTG) to induce protein expression (at 22 °C over 24 h, shaking at 190/min). Protein extraction was performed in 8M urea, 0.1M NaH_2_PO_4_, 0.01M Tris Cl (pH 8) following pelleting of bacteria at 6,000 rpm in a FIBERLite® Rotor (ThermoScientific, F15-8×50cy). The resultant bacterial lysate was centrifuged at 12,000 rpm for 10 min. Pre-cleared supernatant was mixed with Ni-NTA His-Bind Resin (Novagen) (Pajenda et al., 2021, Smits et al., 2021) and incubated under constant rotation (roller mixer SRT6D, Stuart®) for 10 min. The bacterial lysate/resin was loaded onto a Poly-Prep Chromatography Column (Bio-Rad) and drained by unit gravity flow. The column was washed with 8x volumes of 8M urea, 0.1M NaH_2_PO_4_, 0.01M Tris Cl (pH 6.3) also containing 20 mM imidazole. Specifically-bound recombinant protein was eluted in 8 fractions of 300 µl each using elution buffer (8M urea, 0.1M NaH_2_PO_4_, 0.01M Tris Cl (pH 4.5), 250 mM imidazole).

### Immunoblotting of ‘VieVac’ protein with immune sera

Aliquots of the purified protein or its *N* and *S* subregions (Pajenda et al., 2021, Smits et al., 2021) were loaded into individual wells of a 4-20% 10-well SDS-PAGE and run under reducing conditions. The gel was transferred by semidry blotting onto nitrocellulose membranes. The resultant filter was blocked in BM Chemiluminescence Blocking Reagent for 30 min and incubated in 1:100 convalescent serum or 1:1,000 mouse immune serum at room temperature for 2h. Following 2 washes with TPBS (10 min each), goat anti-human IgG (H+L) F(ab’)2 HRP or goat anti-mouse immunoglobuline/HRP was applied and incubated at room temperature under constant shaking for 2 h. Following another 2 washes with TPBS (10 min each), antibody binding was detected by using BM chemiluminescence substrate solutions A and B. Images were recorded with the Fusion FX Vilber Lourmat (Vilber).

### Protein adsorption to Imject™ Alum adjuvant

Next, we tested whether the recombinant protein could adsorb onto Imject™ Alum adjuvant (Hogenesch, 2012). The protein was mixed with the adjuvant in a 1:1 (V/V) ratio, then the potentially formulated vaccine was dialyzed with a Slide-A-Lyzer Dialysis Cassette (Thermo Scientific, MWCO 3500) against PBS at 4°C under constant stirring for 4 h. To demonstrate the adsorption of the immunogen to the adjuvant, the nanostructured vaccine was centrifuged at 12,000 *g* for 4 min. The pre-cleared supernatant and pellet each were treated with Laemmli sample buffer and loaded separately onto a 12% SDS-PAGE gel and run under reducing conditions. The gel was transferred onto nitrocellulose membrans by semi-dry blotting. The filter was then incubated with convalescent sera diluted 1:100 and further developed as described above.

### Mouse immunization

The purified VieVac protein was mixed with Imject™ alum and mice (*n* = 4) were injected intraperitoneally with 20 µg of the prepared immunogen after 5 h of dialysis against PBS. In a second step, the adjuvant AddaVax™ combined with the VieVac protein was dialyzed similarly, then animals were injected with the adjuvant only for the control group (*n* = 5) and with the immunogenic AddaVax/VieVac protein at doses of 10µg, 20µg and 40µg (*n* = 13). In both experiments, a second immunization with the same antigen preparation was administered after 14 days (**Figure 2A)**. Serum was collected 14 and 28 days after the first immunization and tested by ELISA. Gene expression in blood cells and spleen was measured by qPCR at day 28.

### ELISA

Ninety-six-well or 384 well (IgG titer analysis) flat-bottom non-tissue culture treated plates (Falcon) were coated with 100 µl/well of recombinant protein (both *N* and *S*) at a concentration of 1µg/ml. After 1 h of incubation at room-temperature on a plate shaker set to 300 rpm, the antigen-containing fluid was replaced by protein-containing blocking buffer (2x blocking reagent in PBS) for an incubation period of 30 min. Following a brief wash with PBS (200 µl), mouse serum diluted 10^−2^ to 10^−5^ in 100 µl assay buffer was incubated on a plate shaker for 60 min. The ELISA plate was then washed 3x with 350 µl TPBS using an ELISA washing machine (ELX50 Auto Strip Washer, Bio-Tek, Inc.). Antibody binding was determined with goat anti-mouse HRP-conjugate (1:10,000 in assay buffer supplemented with goat serum (2% V/V)) and incubated at room temperature for 60 min. Finally, 100 µl of TMB substrate/chromogen mixture were applied and reacted in the dark for 10 min. Color development was terminated by adding 100 µl of 2M H_2_SO_4_ and read on an ELISA reader (Synergy H1 Hybrid Reader, Bio-Tek, Inc.) at 450 nm. Each sample was processed in technical duplicates. *S* and *N* titers were measured by end-titration using flat-bottomed 384-well Nunc™ plates. ELISA was performed under similar conditions as above, with the reaction volume reduced to 50 µl per well. The reciprocal of the serum dilution was chosen, which gave a 2-fold OD value over the control.

### RNA isolation from blood and spleen

Peripheral blood was subjected to hemolysis using the Erythrocyte Lysis Buffer EL (Qiagen) and RNA was isolated from white blood cells by the MagMAX mirVana Total RNA isolation kit. In brief, lysis buffer mixed with isopropanol was added to each cell pellet and incubated for 3 min. RNA-binding bead mix was added to adsorb RNA onto magnetic beads. Following aspiration of the supernatant, magnetic beads were washed sequentially. TURBO DNase solution was used to prevent genomic DNA contamination, followed by addition of rebinding buffer mixed with isopropanol. Magnetic beads were washed again and after removing the supernatant dried on a rotating platform. Finally, RNA was released by preheated elution buffer and evaluated for its concentration. Tissue fragments of 20-50 mg of mouse spleen were minced in Precellys Ceramic Kit 2.8 mm tubes using the precellys 24 lysis and homogenization device (Bertin Technologies) following the addition of 1000 µl of TRIzol™ Reagent. Subsequently, 200 µl of chloroform were applied and mixed by frequently inverting each tube. Following centrifugation at 12,000 *g* for at 4 °C for 10 min, the aqueous phase was mixed with 500 µl isopropanol. After incubation fat room temperature or 10 min, RNA was pelleted by centrifugation at 12,000 *g* for 15 min. RNA pellets were dried after a wash with 75% ethanol and dissolved in nuclease free water for quantification by a Nano drop device.

### Quantitative PCR

Six hundred ng of RNA were mixed with random primers heated at 65 °C and briefly chilled on ice. dNTPs, RevertAid RT and reaction buffer were added and subjected to first strand synthesis at 55°C for 60 min following primer annealing at 25 °C for 5 min. Enzyme activity was inactivated by heating at 80 °C for 10 min. A 20 µl first strand solution was diluted with 60 µl nuclease-free water and either processed immediately or stored at -80°C. For quantitative PCR, 2 µl of first strand DNA, 1 µl of 10x gene specific ProbeSet from TaqMan, 5 µl of TaqMan 2x universal PCR master mix and 2 µl H_2_O were mixed for each reaction, which all were carried out in duplicate. Reactions were performed on 384 well plates in a QuantStudio 6 Flex machine (Applied Biosystems). At each experiment, 40 cycles were recorded and quantitative data were presented in ΔΔCT mode and showing fold regulation relative to the control. Statistics were calculated using ΔCT values of individual samples.

#### *Ex vivo* T-cell stimulation with N-and S-specific peptides and CD44 evaluation by cell surface flow cytometry

Viable mouse splenocytes were isolated by homogenizing spleen tissues through a 70 µm mesh followed by erythrocyte lysis using the RBC Lysis Buffer (Biolegend) according to the manufacturers’ instructions. Subsequently, splenocytes were seeded in flat bottom 96 well plates in triplicates at 1 × 10^5^ cells per well. For antigen specific stimulation, cells were pulsed with the PepTivator SARS-CoV-2 Prot_S Complete peptide mix (covering the complete sequence of the mature SARS-CoV-2 S protein) or the PepTivator SARS-CoV-2 Prot_N peptide mix (covering the complete sequence of the SARS-CoV-2 N protein; both Miltenyi Biotec). As controls splenocytes were either left unstimulated or activated polyclonally with 12.5 µg/mL phytohemagglutinin (PHA). Cells were then cultured for five days in RPMI supplemented with 10% FCS and 10 µg/mL gentamycin. For analysis of activation, cells were harvested and the respective triplicates pooled. Cells were then stained with anti-mouse CD4 FITC (clone GK1.5), anti-mouse CD8 AlexFluor700 (cloneYTS156.7.7) and anti-mouse CD44 PerCP-Cy5.5 (clone IM7; all Biolegend) in PBS + 0.5% BSA + 0.05%NaN_3_. Expression of the activation marker CD44 was measured on a FACS Canto II flow cytometer (Becton Dickinson) and analyzed using the FlowJo software (TreeStar). For quantification, CD44^high^ cells in the CD4^+^ and CD8^+^ populations from the respective stimulation conditions were corrected against the unstimulated negative control.

### Immunohistochemistry

Free-floating sections were rinsed in PB (0.1M, pH 7.4). Non-specific immunoreactivity was suppressed by incubating the sections in a mixture of 5% normal donkey serum (NDS; Jackson ImmunoResearch) and 0.3% Triton X-100 (Sigma) in PB at room temperature for 2 h. Sections were then exposed (4°C for 3 days) to select combinations of primary antibodies (**Supplementary Table 2**) diluted in PB to which 0.1% NDS and 0.3% Triton X-100 had been added. After extensive rinsing in PB, immunoreactivities were revealed by carbocyanine (Cy) 2-, 3-, or 5-tagged secondary antibodies raised in donkey [1:500 (Jackson ImmunoResearch), at 22–24°C for 2 h]. Glass-mounted sections were coverslipped with Toluol-containing Entellan (Sigma). Sections were inspected and images acquired on a LSM880 confocal laser-scanning microscope (Zeiss) at either 10× or 63× primary magnification with the pinhole set to 0.5 – 0.7 μm. Emission spectra for each dye were limited as follows: Cy2 (505–530 nm), Cy3 (560–610 nm) and Cy5 (650–720 nm). Cell counting was performed in ImageJ. Three sections of the hippocampus (CA1) were analyzed per animal (*n* = 3/experimental group). The stratum pyramidale was excluded from quantification because of its high neuron density.

### Statistical analysis

Adherence to a Gaussian distribution was determined using the Kolmogorov-Smirnov test. Normally-distributed data were provided as means ± s.d. In case of skewed distribution, data were described as medians (25^th^ and 75^th^ percentiles). Qualitative variables were described with counts and percentages and compared using Fisher’s exact test. A two-tailed *p*-value of 0.05 was considered statistically significant. Data were analyzed with SPSS^®^ Statistics (version 21). In histochemical experiments, data were normalized to a surface area of 1 mm^2^ and expressed as means ± s.e.m, followed by one-way ANOVA in GraphPad Prism.

## Results

### Study rationale

Earlier work in infected individuals showed abundant antibody production against parts of the nucleocapsid, while neutralizing antibodies were shown to recognize regions in the RBD (and also NTD) of the spike protein (Sterlin, Mathian et al., 2021, Wu et al., 2020). These findings prompted us to design a fusion protein including immunodominant regions of bot the *S* and *N* proteins. (**Fig. 1A**) Fig. S1 shows the resultant divalent recombinant nucleotide and protein sequences, which we sought to test for their immunogenicity.

**Figure 1:**
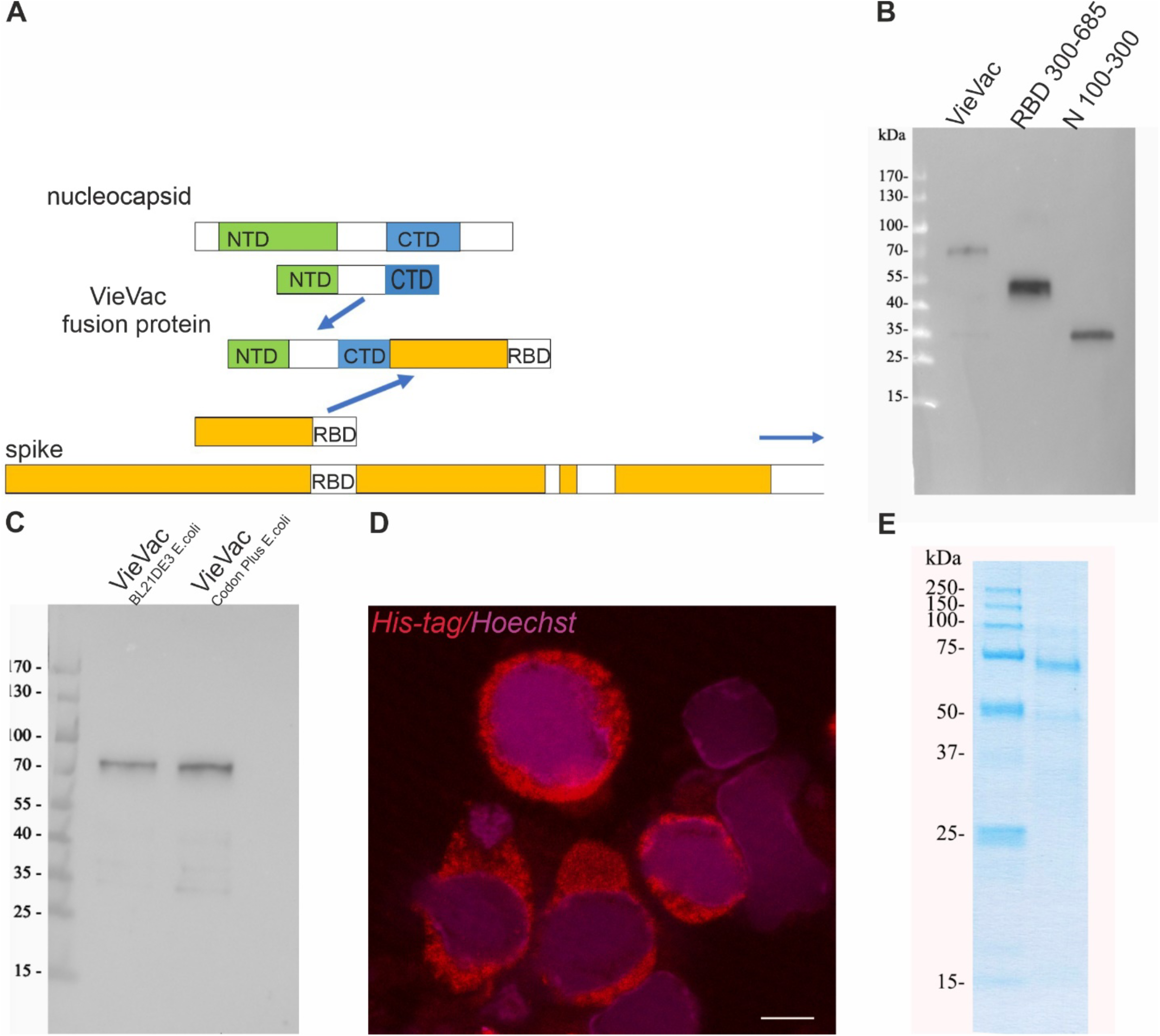
Expression construct and biochemical characterization of purified proteins. **A:** Scheme of fusion protein design. The immunodominant region *N*100-300 aa of the nucleocapsid and *S*300-685 aa of the spike protein were fused and the product (termed ‘VieVac’) was engineered into either the pET-30a *E. coli* expression vector or a baculovirus. **B:** Protein immunoblot of *E. coli*-produced ‘VieVAc’ fusion protein and its constituents using convalescent serum. The fusion protein (‘VieVac’, lane 1), *S*300-685 aa (lane 2) and *N*100-300 aa of the nucleocapsid protein (lane 3). Molecular weight marker is shown to the left. **C:** Protein immunoblot of *E. coli*-produced fusion proteins ‘VieVac’ using mouse immune serum. The fusion proteins were purified out of *E. coli* lysate through their His-Tag. VieVac produced in *E. coli* BL21DE3 (lane 1), ‘VieVac’ produced in *E. coli* BL21Codon Plus (lane 2). **D:** Immunofluorescence staining of Hi5 cells using anti-His.H8 mAb to detect ‘VieVac’ (red) in cells infected with baculovirus. *Scale bar* = 10 µm. **E:** Protein staining of insect cell-produced fusion protein ‘VieVac’. The fusion protein was purified through its His-Tag out of Hi5 lysate results, with the full-size protein migrating at 72 kDa with only minor degradation products (lane 2). Molecular weight markers are shown to the left (lane 1).

### Protein production, characterization and purification

First, we produced and purified the fusion protein (termed ‘VieVac’) in *E. coli*. The recombinant protein containing a 6-His-tag was successfully purified by 8M urea. As shown in **Fig. 1B,C**, the recombinant protein migrated at ∼70 kDa, which is at the calculated cumulative molecular weight of the *N*100-300 aa (lane 3 in **Fig. 1B**,) and *S*300-685(lane 2 in **Fig. 1B**). In addition to the entire ‘VieVac’ protein, a truncated fragment co-purified in most experiments due to a premature translation stop (lane 2 in **Fig. 1C**).

To evaluate whether the ‘VieVac’ protein would adsorb to AOH and AlPO_4_-based adjuvant (Imject™ Alum), immunoblotting of formulated ‘VieVac’ was performed after pelleting of the nanoparticulate structures by centrifugation. This revealed that the protein was recovered in its entirety in the particle-containing pellet (**Fig S2**, lane 3).

In a second attempt, ‘VieVac’ production was performed in eukaryotic cells. This was motivated by the fact that the RBD is structurally composed of a twisted five-stranded antiparallel β-fold, with strands and loops connecting the β-strands, and is kept in its configuration by 4 disulfid bonds between 8 cysteins. Since this delicate type of folding develops better in eukaryotic cells, the recombinant gene fusion was re-engineered into the baculovirus system with the non-truncated protein successfully produced in Hi5 cells as demonstrated by immunofluorescence labelling 72 h after infection (**Fig. 1D**). This protein could be purified through its N-terminal His-tag in its full length. Only minor degredation products co-purifed on Ni-NTA colums (**Fig. 1E**).

### Imject™ Alum/’VieVac’ induces IgGs to both nucleocapsid and RBD

To reduce the required amount of the antigen, the recombinant fusion protein was adsorbed onto an AlOH and AlPO_4_-based adjuvant. First, mice (*n* = 4) were challenged with this immunogen in a standard regimen (Fig. 2A) (Wagner, Gessl et al., 1996). Immunogen-challenged mice developed IgG by 14 days (nucleocapsid titer 10^−2^ - 5×10^−2^ and RBD 10^−2^ - 10^−3^) after the first dose, which increased to a significant IgG response to both antigens 28 days after the initial injection (nucleocapsid end-titer 10^−5^ and RBD end-titer 5×10^−4^ - 10^−5^) (**Fig. 2B-F**). None of the mice showed any adverse side effect, e.g. weight loss or neurological complications (*data not shown*). Mouse #1 had a shallow IgG response towards the nucleocapsid after the first booster injection (end-titer 10^−2^) (**Fig. 2B**). When receiving a second booster, this mouse also responded to the nucleocapsid when tested later (**Fig. 2D**) with an end-titer of 10^−5^ (**Fig. 2E**).

**Figure 2:**
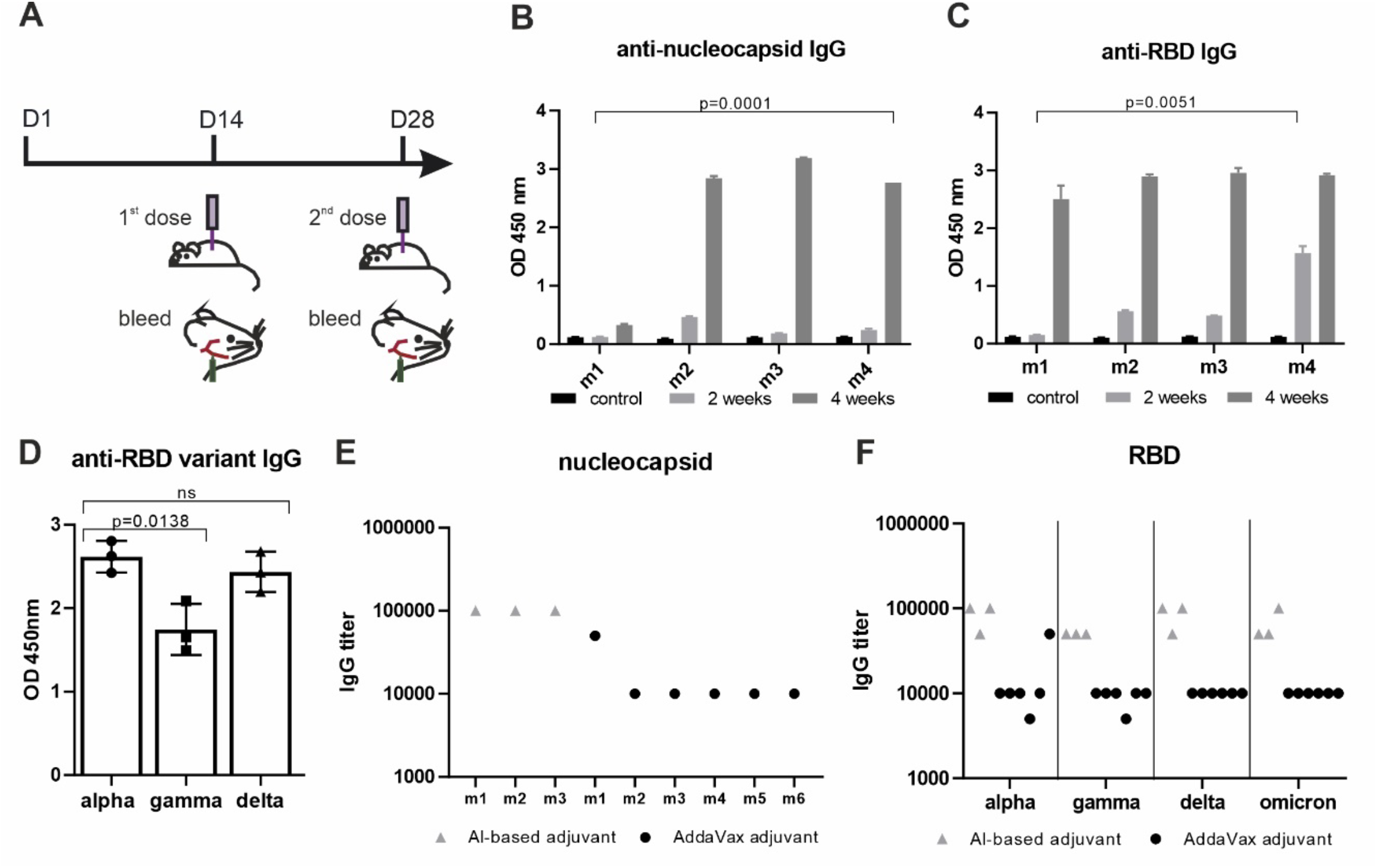
‘VieVac’ produces prolonged immune responses against all known virus strains. **A:** Adult C57BL/6J mice (*n* = 4) were immunized with 20 µg Imject™ Alum /VieVac (i.p.) with a booster injection carried out 14 days later. Blood sampling was performed 1 h before immunization and on day 28 after the first immunization. **B:** Anti-nucleocapsid and **C:** anti-spike IgG response measured by ELISA upon administration of Imject™ Alum/’VieVac’ in 4 mice (m1, m2, m3, m4) 2 weeks (grey) or 4 weeks after the first injection (dark gray), as compared to controls (black). **D:** Anti-RBD IgG cross-reactivity in mice (*n* = 3) against the alpha variant, relative to the gamma and delta variants treated with the prime booster mode of Imject™ Alum /’VieVac’ 90 days prior. **E:** Anti-nucleocapsid IgG end-titer in Imject™ Alum /VieVac (gray) and AddaVax™/’VieVac’ (black) prime booster immunized mice. **F:** Anti-RBD IgG end-titer (alpha, gamma, delta, omicron variants) in Imject™ Alum /’VieVac’ (gray) and AddaVax™/’VieVac’ (black) prime booster immunized mice.

In order to investigate the durability of the IgG immunogen response, mice were kept under normal conditions and tested after 90 days for the mutant RBD of SARS-CoV-2 variants termed gamma (K417T and E484K) and delta (L452R and T478K). This revealed immune cross-reactivity of alpha RBD with the delta variant and significantly (*p* = 0.0138, 19%) reduced cross-reactivity with the gamma variant (**Fig. 2D**). Mouse #1 having received two booster injections showed similar reactivity towards both the alpha and delta variants.

### AddaVax™ with ‘VieVac’ primes IgG production to both nucleocapsid and RBD

In a second study, ‘VieVac’ generated in a eukaryotic expression system was formulated with AddaVax™, an MF59 compatible adjuvant, which is approved in Europe and has been used in influenza vaccines (Mbow, De Gregorio et al., 2010) and beyond (Chappell, Mordant et al., 2021). Mice (*n* = 13) were again challenged with 3 different doses of recombinant divalent antigen mixed with AddaVax™ according to the benchmarked immunization strategy used above (see **Fig. 2A)**. Ten out of thirteen recipients produced IgG against both proteins (nucleocapsid titer: 10^−2^ - 10^−3^ and RBD titer :10^−2^ - 10^−3^) after 14 days. Control mice received AddaVax™ only (*n* = 5). Fourteen days after a booster injection, 9 out of the 13 test mice showed a further increase in their IgG responses to both antigens (nucleocapsid titer: 10^−4^ - 5×10^−5^ and RBD titer: 5×10^−3^ - 10^−5^) **(Fig. 2E-F and Fig. 3A_1_-A_2_**).

**Figure 3:**
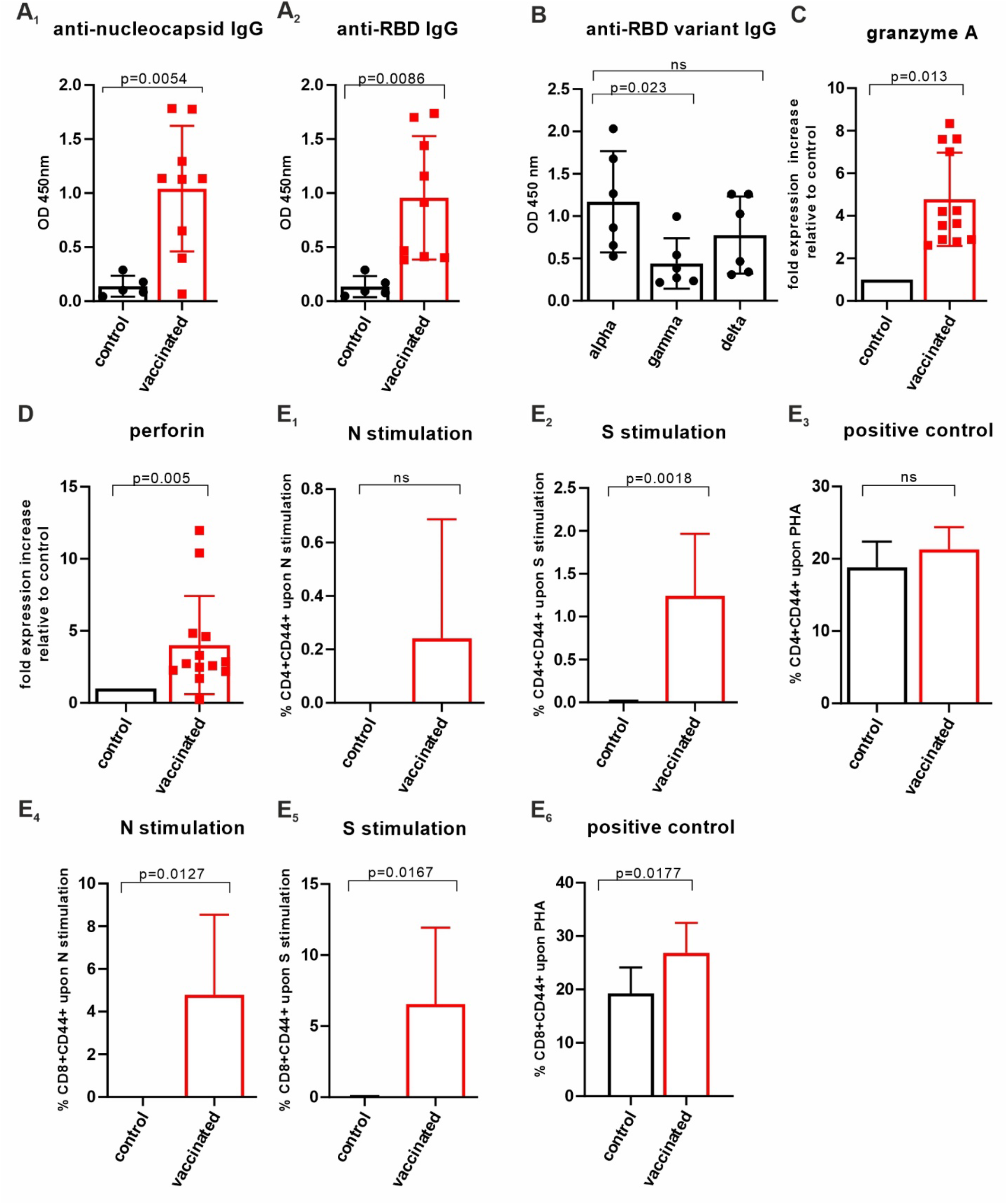
Assessment of cellular immunity upon AddaVax™/’VieVac’ administration. **A**_**1**_**-A**_**2**_: AddaVax™/’VieVac’ immunisation in prime-booster mode and IgG responses in 9 out of 13 mice. Control mice (n = 5) received AddaVax™ only, test animals received AddaVax™/’VieVac’ containing 10 to 40 µg protein. **B:** Cross-reactivity against SARS-CoV-2 RBD variants, alpha *vs*. gamma and delta. Sera of *n* = 6 mice prime-booster injected with AddaVax™/’VieVac’ were tested for IgGs recognizing alpha, as compared to gamma and delta versions of the RBD. **C-D:** Effector molecule gene expression in peripheral blood. Granzyme A (C) and perforin (D) expression in peripheral lymphocytes was evaluated by qPCR using a TaqMan probe set. Relative fold expression is indicated in 12 AddaVax™/’VieVac’ prime-booster injected mice in relation to the mean of *n* = 5 control mice that had received AddaVax™ adjuvant only. **E**_**1**_**-E**_**6**_: CD4^+^ and CD8^+^ T-cell responses in *ex vivo* stimulated spleen cells. CD4^+^ T-cells (E_1_-E_3_) were stimulated *ex vivo* with N-specific (E_1_) and S-specific peptides (E_2_) or PHG (E_3_). Control mice (*n* = 5, adjuvant only) were compared to vaccinated mice (*n* = 13, AddaVax™/’VieVac’). CD8^+^ T-cells (E_4_-E_6_) were stimulated *ex vivo* with N-specific (E_4_) and S-specific peptides (E_5_) of PHA (E_6_). Control mice (*n* = 5, adjuvant only) were compared to vaccinated mice (*n* = 13, AddaVax™/’VieVac’).

Next, we tested mouse sera (*n* = 6) for individual IgG cross-reactivity against the virus variants gamma and delta. As compared to alpha, each serum IgG content was tested 14 days after booster injection for its reactivity against the mutant RBD (**Fig. 3B)**. The immune response (per IgG) varied amongst the mice but was reduced on average by 63% (*p* < 0.023) towards the gamma variant and 43% (non-significant) towards the delta variant, as compared to the alpha variant when measured at a titer of 10^−2^. However, similar end-titers were found for alpha, gamma-delta, and omicron variants, ranging from 5×10^−3^ to 10^−4^, with the exception of one mouse in which the end-titer was an order of magnitude lower when alpha was compared with gamma, delta and omicron **(Fig. 2F)**. These data show that our #VieVac’s trategy is equally efficient against all prevalent SARS-CoV-2 strains.

### Insect cell-produced AddaVax™/VieVac generates effector cell immunoreactivity

Next, we addressed the effector potential of the constructs injected by analyzing cytotoxic lymphocytes (NK-T-cells), which have been suggested as the most important determinants of cellular anti-SARS-CoV-2 immunity. In peripheral blood, the AddaVax™/VieVac immunized group (*n* = 13) showed a 4.6-fold (*p* < 0.013) granzyme A and a 4-fold (*p* < 0.005) perforin mRNA increase, as compared to the AddaVax only control group (*n* = 4). (**Fig. 3C,D**). In order to investigate changes in the lymphocyte populations in the spleen, 20-25 mg of spleen tissue was homogenized, and RNA extracted. When analyzed for changes in target gene expression, changes did not reach the level of significance, even though an up-regulation in granzyme A expression was seen (**Table S1**).

### CD4^+^ and CD8^+^ T cell responses evaluated by CD44 cell surface upregulation in *ex vivo* stimulated spleen cells of AddaVax™/VieVac challenged mice

N-specific and S-specific T-cell responses were measured following 5 days of *ex vivo* spleenocyte stimulation using T-cell-specific peptides of SARS-CoV-2 *N* and *S* protein. Spleen T-cells were obtained 14 days after booster injection from mice injected with AddaVax™/VieVac (*n* = 13) as test group and adjuvant (AddaVax™) only (*n* = 5) as control group. The number of CD4^+^ T-cells expressing CD44 upon stimulation by N-specific peptides was elevated in 7 mice out of 13 **(Fig. 3E_1_**) but did not reach statistical significance when compared to the control group. Stimulation with *S*-specific peptides resulted in a significant increase in CD44 in the test group compared with the control group (*p* = 0.0018; **Fig. 3E_2_**). The extent of activated CD8^+^ T-cells after stimulation with *N*-specific peptides (*n* = 13) and *S*-specific peptides (*n* = 13) was significant relative to the control group (AddaVax™ only, *n* = 5; *p* = 0.0127 and *p* = 0.0167 respectively. **(Fig. 3E_4_-E_5_**). Phytohemagglutinin (PHA) stimulation was used as a positive control to illustrate the spectrum of CD44 response to a nonspecific T-cell stimulant (**Fig. 3E_3_, E_6_**). *S*-specific stimulated CD8^+^ T-cells showed significantly higher stimulation by PHA in terms of CD44 upregulation compared to control (*p* = 0.0177) **(Fig. 3E_6_**). This suggests that a higher state of alertness to non-specific stimuli is induced within the CD8^+^ T-cells after S-specific peptide exposure.

### Lack of adverse neuropathology in Addavax™/VieVac-injected mice

Neurological side effects have been observed following vaccination with authorized viral vector-based vaccines against SARS-CoV-2 (Chun, Park et al., 2022, Greinacher et al., 2021, Sharifian-Dorche, Bahmanyar et al., 2021, Zuhorn et al., 2021). Particularly, cerebrovascular venous and sinus thrombosis (CVST) is a subtype of stroke in which blood clots form and obstruct blood flow in the brain’s vascular system (Stam, 2005). These conditions combined with a COVID vaccine-related thrombocytopenia have been termed vaccine-induced immune thrombotic thrombocytopenia (VITT) (Greinacher et al., 2021, Schultz, Sørvoll et al., 2021) and this has become one of the most important research areas of our time.

No differences were detected in endothelial morphology or blood vessel density/distribution, between experimental groups **(Fig. 4A-D**,) (Härtig, Reichenbach et al., 2009). Reduced capillary staining, the presence of plaques, and ramified (activated) microglia were not detected either. Likewise, the distribution, density or morphology of microglia and astrocytes **(Fig. S6A-D)** also remained unchanged **(Fig. 4A-D)** (Bignami, Eng et al., 1972, Sasaki, Ohsawa et al., 2001). No differences were observed in the cell number of neurons in selected brain areas either **(Fig. 4E-F and S5)**. These data support the notion that ‘VieVac’, at least in preclinical models, is unlikely to cause adverse neuropathological side-effects.

**Figure 4:**
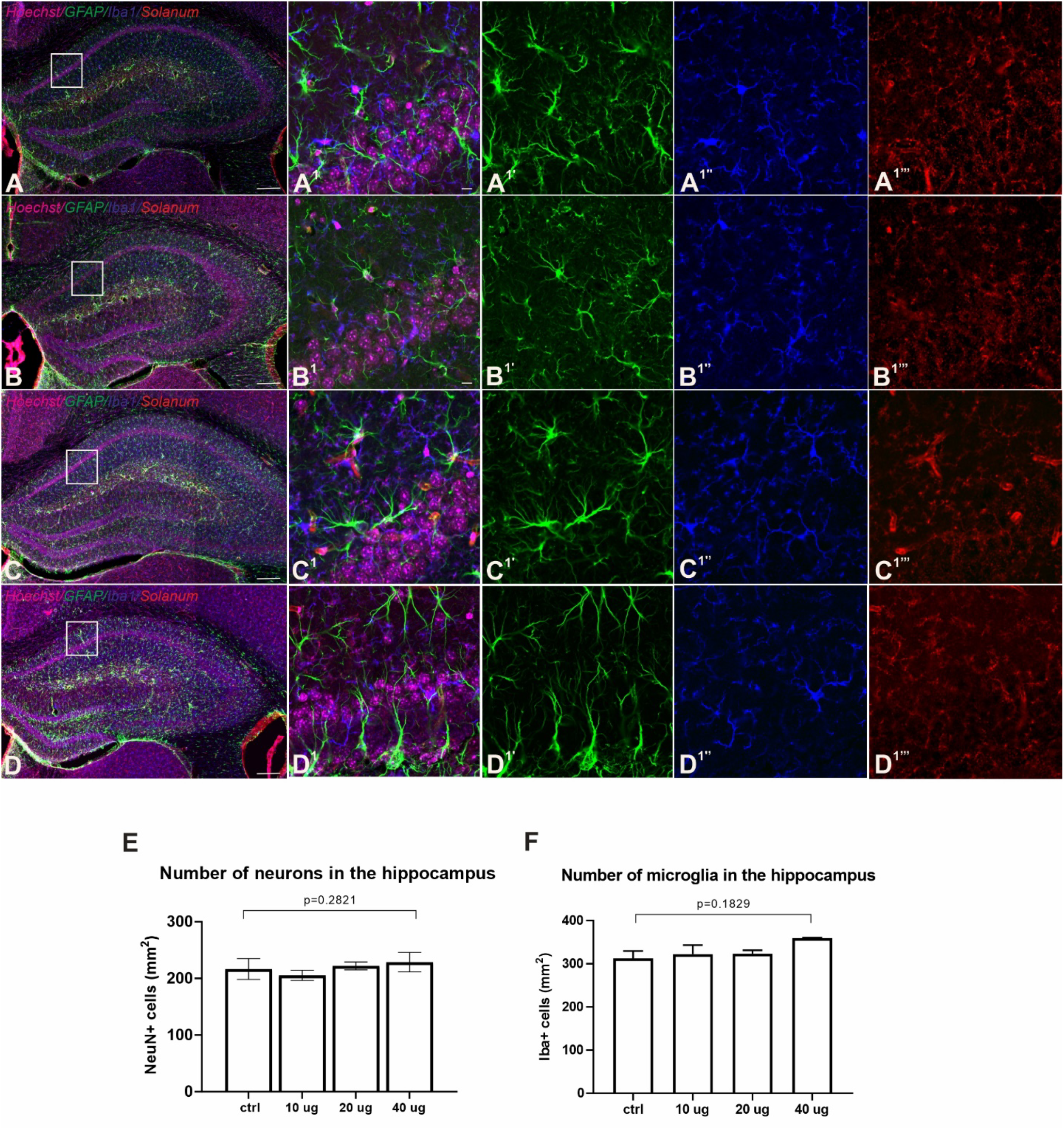
Lack of adverse side-effects in the nervous system of immunized mice. **A-D:** Fluorescence immunohistochemistry showed no accumulation in GFAP (green, astrocytes, A1’- D1’), Iba1 (blue, microglia, A1’’-D1’’) and *Solanum tuberosum* lectin (red, vasculature, A1’’’-D1’’’) distribution between control (A) and Addavax™/’VieVac’-injected animals (B: 10 µg, C: 20 µg, D: 40 µg). **E-F**: Quantitative analysis of neuronal numbers (NeuN; E) and Iba1-positive microglia (F) in the hippocampus (CA1 subfield) showed no difference between the control and vaccinated groups. Scale bars = 200 µm (A-D), 10 µm (A^1^-D^1’’’^).

## Discussion

This study provides a proof-of-concept approach for the generation of divalent (or even polyvalent) protein backbones for the development of protein vaccines against SARS-CoV-2 taking advantage of the coincident presence of the most immunogenic viral regions hinged by flexible linkers to maintain ternary and quaternary structures. The construct design incorporates a rapidly mutating region (RBD) and a constitutive region (nucleocapsid), thus overcoming strain-specific hindrances in immunogenicity as increasingly seen for linearized mRNA vaccines.

The RBD region is the backbone of all vaccination strategies presently available, even if its inferiority upon rapidly mutating SARS-CoV2 is already apparent. Since conformation-specific antibody responses may be important in immune defense against SARS-CoV2, recombinant proteins generated in eukaryotic systems are the tools of choice. This is significant because the RBD domain has a twisted five-stranded antiparallel ß-fold with strands and loops connecting its ß-strands. This fold could be resolved by crystallography of the RBD produced in Hi5 insect cells (Lan, Ge et al., 2020). There are 9 cysteines in this region, 8 of which are involved in disulfide bond formation in order to generate the ACE-2 binding structure. Here, we show that insect bioreactor systems could also be amenable to producing vaccine backbones. Particularly, our molecular design incorporates 8 cysteines from the RBD region (300-685 aa), thus likely stabilizing the ternary structure of the protein to increase its recognition by the host’s immune system, a feature likely contributing to a near-maximal immune response already at the second booster stage. In contrast, relatively limited emphasis has been directed towards the well-transcribed nucleocapsid protein in presently pursued SARS-CoV2 vaccine strategies in Europe and beyond, even if the nucleocapsid of other coronaviruses was earlier recognized for its immunogenicity (Boots, Kusters et al., 1991). This lack of interest is surprising since 11% of human CD4+ T-cells and 12% of CD8+ T-cells recognize the nucleocapsid in individuals with SARS-CoV2 infection (Grifoni, Weiskopf et al., 2020), supporting its immunogenic potential to generate potent immunoprotection by expanding surveillance to cellular branches of defense in humans.

A homologue of an EDA-approved adjuvant was used to increase efficacy, which was in terms of T-cell response near-complete after a second booster, alike by others (Chappell et al., 2021), and reached 100% upon a third inoculation in a staggered primer-booster regimen. The fact that high immunogenicity is detected even after 90 days in a mouse immunization model with near-equivalent efficacy against the alpha, delta and omicron strains when measuring the end-titer gives confidence in the correctness of this design strategy. Immunological analysis of lymphocytes harvested from the spleen of immunized animals showed a significant T-cell response as indicated by the significant upregulation of granzyme A and perforin, cytotoxic granule effector molecules, in peripheral lymphocytes. An increased expression of CD44 upon N-and S-specific peptide stimulation ex vivo demonstrated the generation of T-cells capable of executing cellular protection and production of interleucins (Tian, Patel et al., 2021).

Given the brevity of time for our compressed proof-of-concept workflow, we recognize that data on the generation of neutralizing antibodies, whose presence is assigned to immunity against the RBD (‘spike’) protein, is as yet lacking. However, and equally importantly, cellular immunity through T-cell responses is recognized as an essential means of protection. Accordingly, the adoptive transfer of T-cells into immunodeficient mice led to rapid recovery after the transfer of SARS-CoV-specific effector cells (Zhao, Zhao et al., 2009). Similar results are seen in ferrets, in which neutralizing antibodies do not fully protect against SARS-CoV2-induced disease. Instead, nasal immunity, which is reliant on T-cell responses together with the presence of antibodies, seems to carry optimal protection (See, Petric et al., 2008). Notably, nucleocapsid-related B-cell immunity has also been shown earlier (Guo et al., 2020, Pajenda et al., 2021, Shrock, Fujimura et al., 2020, Smits et al., 2021, Wang et al., 2020), and even served as a diagnostic tool because the detection of nucleocapsid-specific IgG in conjunction with RBD-specific IgG differentiated SARS-CoV2 exposure from a pure mRNA-based vaccine response. Therefore, the pronounced B-cell responses shown here support the hypothesis that both viral protein fragments of our divalent construct can provoke significant cellular immunity. Considering that the immunogenic peptides are native viral sequences (and neither forward nor reverse-engineered fragments that could curb immunogenicity), we view the B- and T-cell responses described here as minimally required yet sufficient experimental indices of our molecular design strategy to trigger significant protection against SARS-CoV2.

In conclusion, we propose that the above molecular design together with biological data on the efficacy of the administered divalent recombinant protein outline a rational approach to generate an efficient protein vaccine pipeline (which we term as a prototypic “VieVac” vaccine), which can be built on both eukaryotic and prokaryotic bioreactors for scaleable production. The bivalent protein/adjuvant-based immunization is effective in eliciting humoral and cellular immunoresponses upon application in a prime-booster mode. Given that the repeated administration of “VieVac” evoked no side effects in the nervous system, we emphasize the safety of a protein-based vaccine.

## Acknowledgments

We want to thank Reingard Grabherr and Miriam Klausberger for instructions on insect cell culture, Monika Aiad for technical support, Angelika Freudenthaler and Sabina Baumgartner-Parzer for laboratory management. This work was in part supported by the European Research Council (ERC-2020-101021016; T. Ha.).

## Supplementary information

**Supplementary Figure 1.**
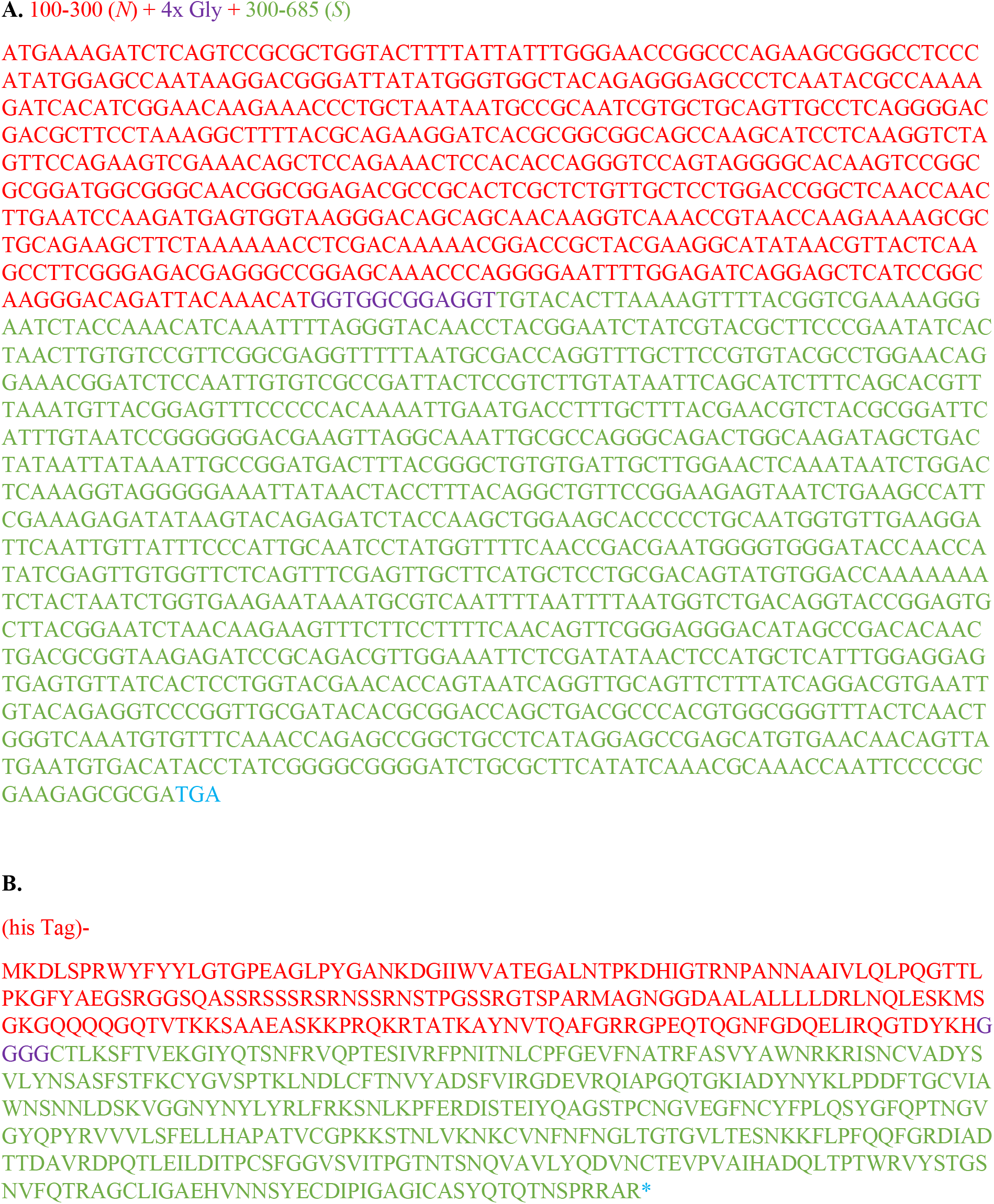
‘VieVa’ coding (A) and translated sequence (B)

**Supplementary Figure 2.**
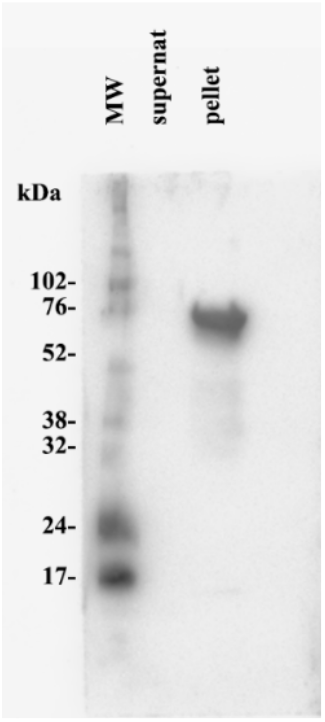
Adsorption of recombinant ‘VieVac’ protein to Imject™ Alum and immunoblotting with convalescent serum. Equal volumes of fusion protein solution and Imject™ Alum were incubated at room temperature for 5 min under constant rotation. An aliquot (20 µl) was centrifuged at 12,000 *g*. Pre-cleared supernatant (lane 2) did not contain any ‘VieVac’ protein but all was adsorbed onto the pelleted particles of Imject™ Alum (lane 3). Molecular weight (MW) markers are in lane 1.

**Supplementary Figure 3.**
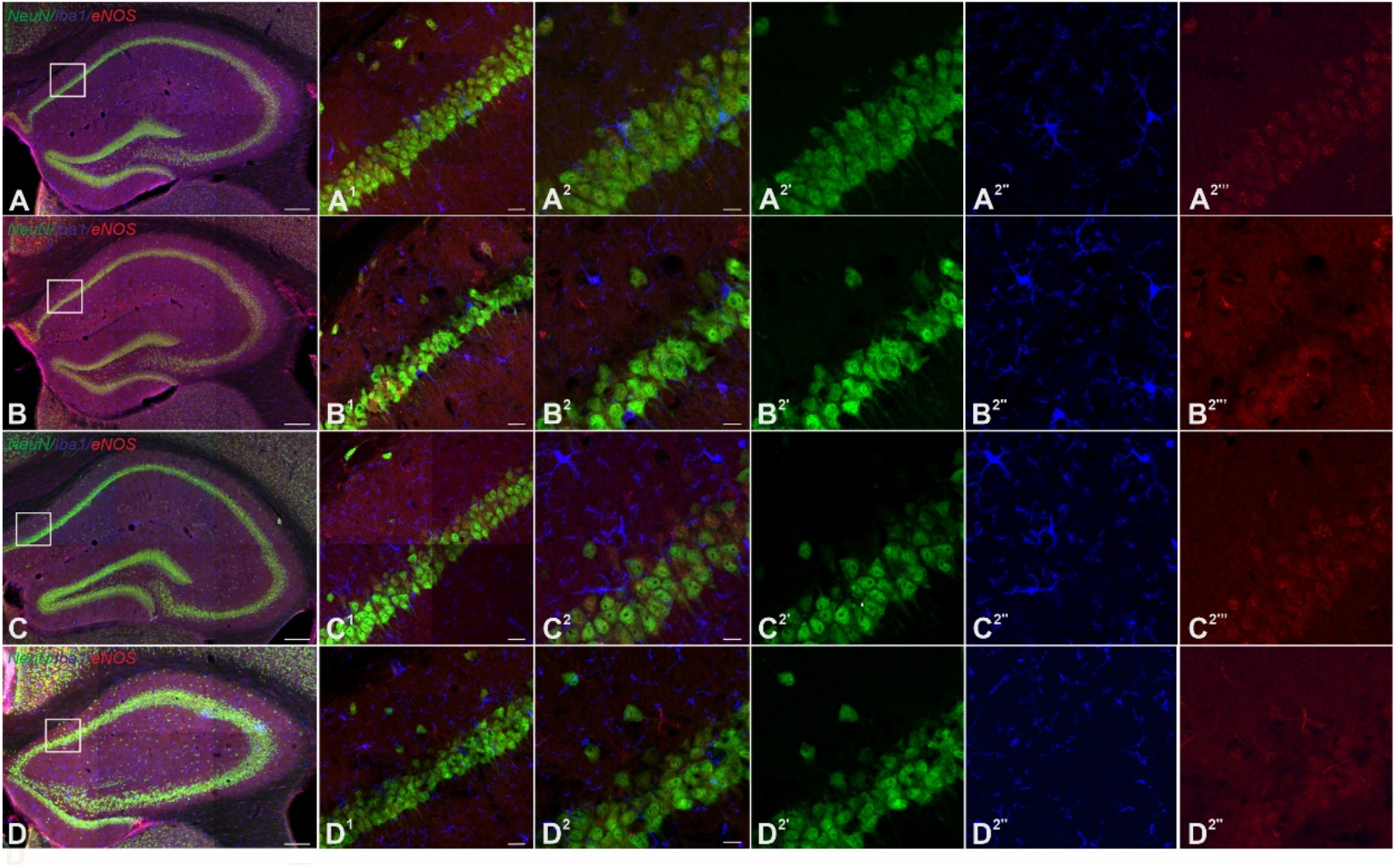
Multiple immunofluorescence labelling confirmed the lack of adverse effects on the density and morphology of neurons and microglia in the hippocampus of immunized mouse brain. NeuN (green, neurons, A2’-D2’), Iba1 (blue, microglia, A2’’-D2’’) and eNOS immunoreactivities (red, vasculature, A2’’’-D2’’’) were assessed in control (A) and Addavax™/’VieVac’-injected animals (B: 10 µg, C: 20 µg, D: 40 µg). *Scale bars* = 200 µm (A-D), 20 µm (A^1^-D^1^), 10 µm (A^2^-D^2’’^).

**Supplementary Figure 4.**
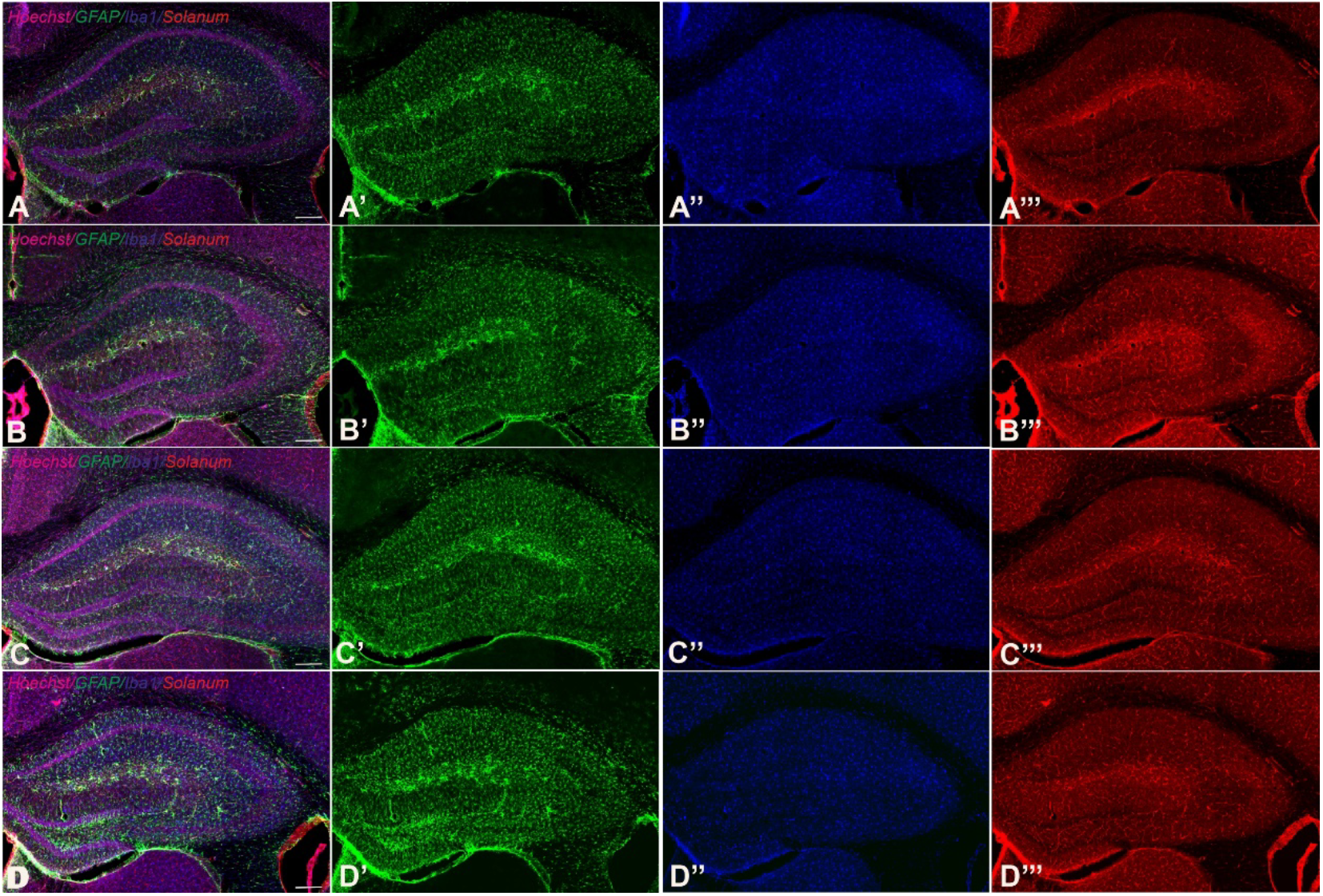
Multiple immunofluorescence labelling confirms the lack of inflammation in immunized mouse brain. GFAP (green, astroglia, A2’-D2’), Iba1 (blue, microglia, A2’’-D2’’) and eNOS immunoreactivities (red, vasculature, A2’’’-D2’’’) were assessed in control (A) and Addavax™/’VieVac’-injected animals (B: 10 µg, C: 20 µg, D: 40 µg). *Scale bars* = 200 µm.

**Supplementary Table 1.** Gene expression in spleen measured by qPCR in control (AddaVax adjuvant only, *n* = 4) *vs*. immunized mice (Addavax™/’VieVac’, *n* = 13) 28 days after the initial injection. No statistical difference in gene expression was found. A trend towards up-regulation in granzyme A was seen in the test group.

| **Target Name** | **probe set, TaqMan** | **P value** |
| --- | --- | --- |
| VEGFA | Mm01281447_m1 | 0,5953 |
| CD19 | Mm00515420_m1 | 0,7370 |
| CD3e | Mm1179194_m1 | 0,7627 |
| CD8a | Mm01182108_m1 | 0,6222 |
| Granzyme A | Mm01304452_m1 | **0,3351** |
| TGFb1 | Mm03024053_m1 | 0,8565 |
| IFNg | Mm01168134_m1 | 0,8044 |
| Perforin | Mm00812512_m1 | 0,5776 |

**Supplementary Table 2.** Antibodies and their use for biochemistry and histochemistry.

| **Marker** | **Company** | **Host** | **IH dilution** | **Catalogue no.** |
| --- | --- | --- | --- | --- |
| eNOS | Invitrogen | rabbit | 1:300 | PA3-031A |
| GFAP | Synaptic System | guinea pig | 1:1000 | 173004 |
| Iba1 | Abcam | goat | 1:500 | Ab5076 |
| NeuN | EMD Millipore | chicken | 1:1000 | ABN91 |
| Solanum tuberosum<br>biotinylated lectin | Vector<br>Laboratories | - | 10 µg/µl | B-1165-2 |
| Hoechst 33,342 | Sigma | - | 1:10.000 | 23491-52-3 |
| His tag HIS.H8 | Millipore | mouse | 1:500 | 05-949 |
| anti-human IgG (H+L)<br>F(ab')<sub>2</sub> HRP | Millipore | goat | 1:20.000 | AQ112P |
| anti-mouse<br>immunoglobulin/HRP | Dako | goat | 1:10.000 | P0447 |

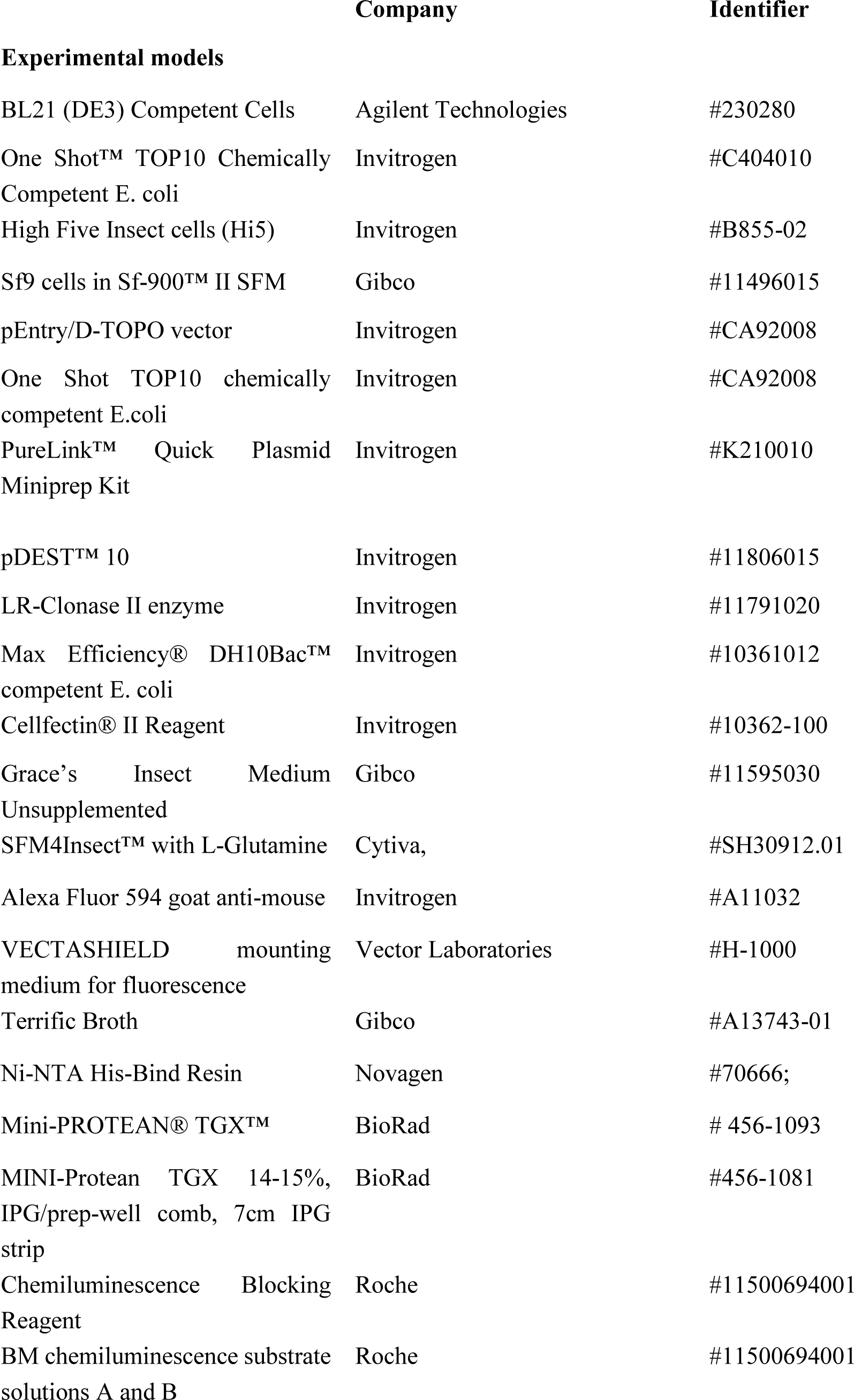
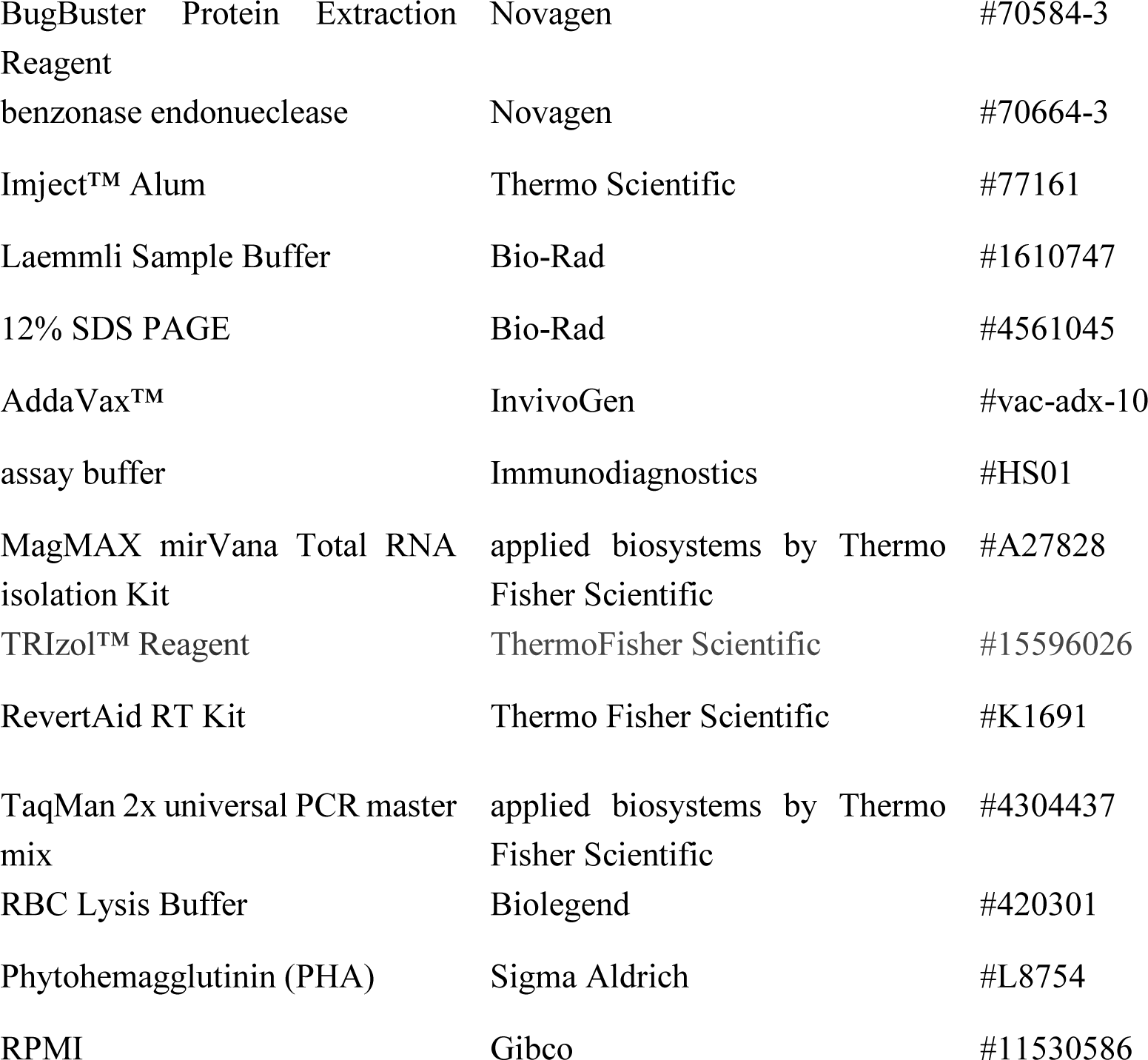

